## Supplementary material for "Global FAW population genomic signature supports complex introduction events across the Old World": Suppl. Fig.

**(1A)**

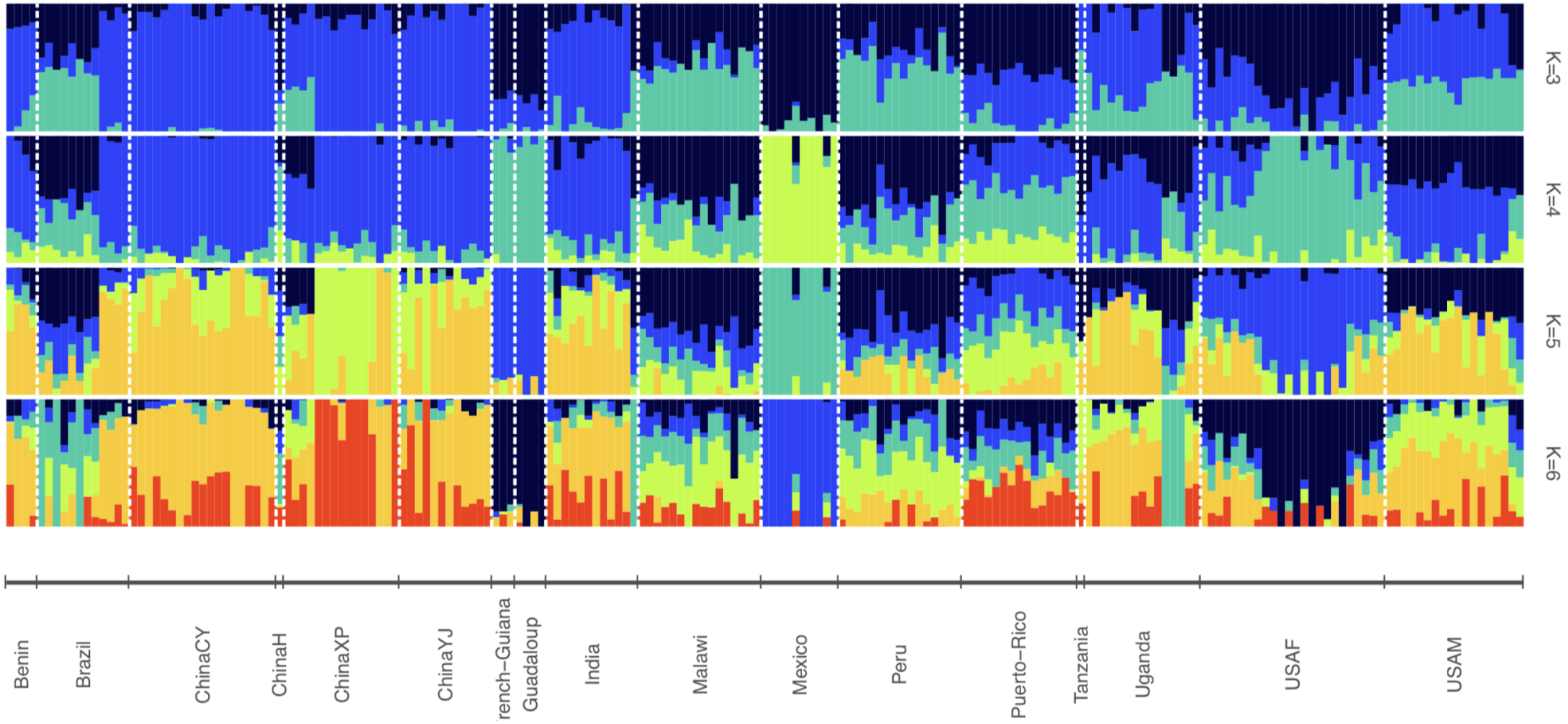


**Suppl. Fig. 1:** (**1A**) Admixture analysis based on 870 unlinked and neutral genome-wide SNPs for FAW populations from both native (North, Central, South Americas) and invasive (Africa, Asia) ranges. The Mexican population represents the only New World native FAW population with most individuals having non-admixed genome patterns, and where all individuals also have the corn mitogenome haplotype. One Mexican individual also have the rice *Tpi* signature (1B).

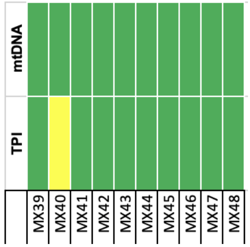


**(1B)** Mitochondrial DNA genome and *Tpi* gene characterisation of Mexican FAW.

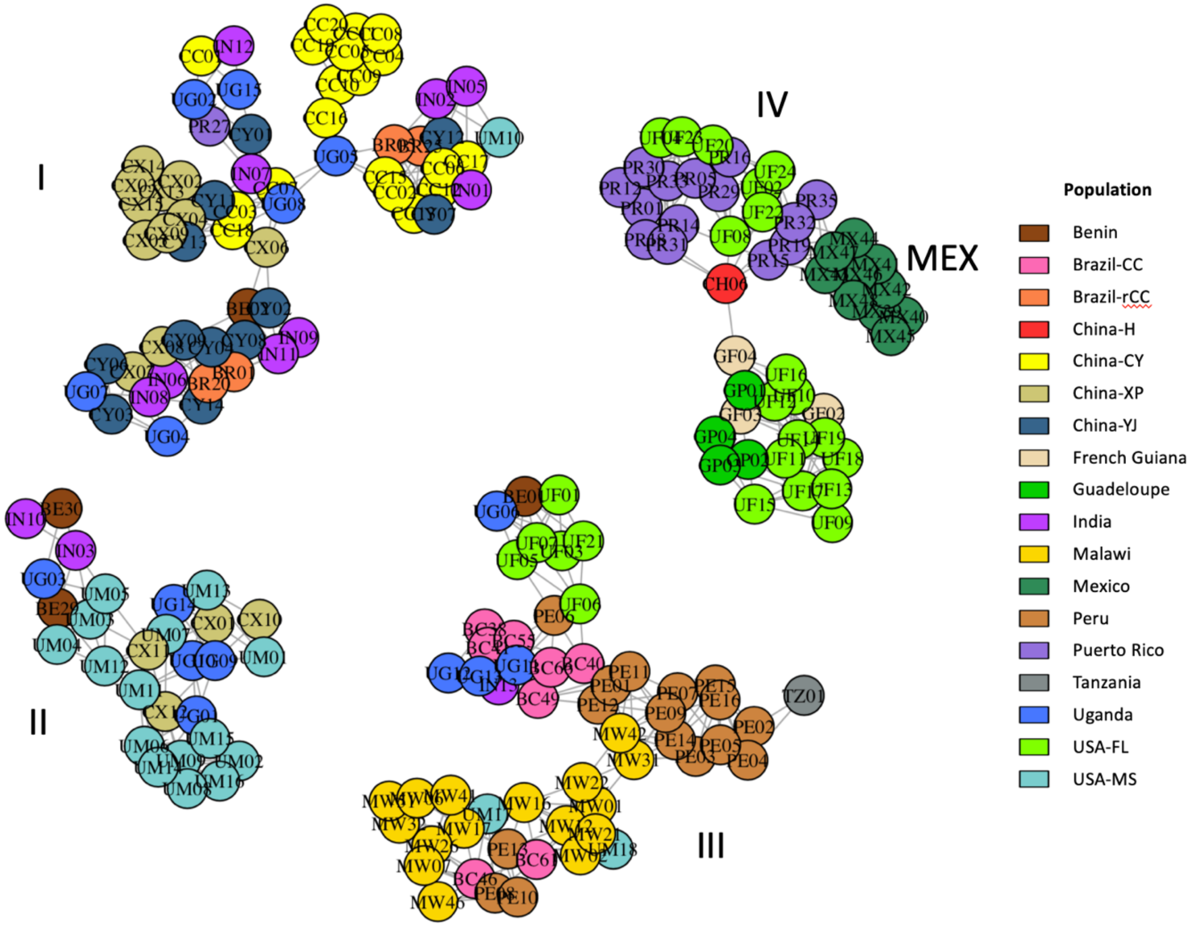


**Suppl. Fig. 2:** Maximum Likelihood (ML) network showing individuals belonging to the populations as given in the legend. The network was drawn using the plotAdmixture function in the R package NetView [1,2], and is based on a ML distance matrix calculated from the IQ-Tree shown in Fig. 3. using the R package ape [3]. The four major clusters, I – IV, correspond to those shown in the tree. Individuals are identified by country codes as follows: China XP (CX), China YJ (CY), China CY (CC), China CH06 (CH), India (IN), Uganda (UG), Tanzania (TZ), Malawi (MW), Benin (BE), Brazil CC (BC), Brazil rCC (BR), Peru (PE), French Guiana (GF), Mexico (MX), Guadeloupe (GP), Puerto Rico (PR), USA-Florida (UF), and USA-Mississippi (UM). See Supplementary Table 1 for complete information about the individuals.
