## Supplementary material for "Global FAW population genomic signature supports complex introduction events across the Old World": Suppl. Table

**Supplemental Table 1:** *Spodoptera frugiperda* samples used in this study including sample and population codes, host preferences based on mitochondrial DNA COI partial gene characterisation, and GenBank accession numbers for assembled mitogenomes.

| Sample | continent | Country | location | Popcode | Sample-ID | mtGenome | from | GenBank Acc. No. |
|---|---|---|---|---|---|---|---|---|
| CH06 | Asia | China | Yunnan | CH | HIA006 | corn | CSRIO/DAWE | MT897262 |
| CY01 | Asia | China | Yuanjiang | CY | YJ1 | rice | NJAU | MT897263 |
| CY02 | Asia | China | Yuanjiang | CY | YJ2 | rice | NJAU | MT897264 |
| CY03 | Asia | China | Yuanjiang | CY | YJ3 | rice | NJAU | MT897265 |
| CY04 | Asia | China | Yuanjiang | CY | YJ4 | rice | NJAU | MT897266 |
| CY06 | Asia | China | Yuanjiang | CY | YJ6 | corn | NJAU | MT897267 |
| CY07 | Asia | China | Yuanjiang | CY | YJ7 | rice | NJAU | MT897268 |
| CY08 | Asia | China | Yuanjiang | CY | YJ8 | rice | NJAU | MT897269 |
| CY09 | Asia | China | Yuanjiang | CY | YJ9 | rice | NJAU | MT897270 |
| CY11 | Asia | China | Yuanjiang | CY | YJ11 | rice | NJAU | MT897271 |
| CY12 | Asia | China | Yuanjiang | CY | YJ12 | rice | NJAU | MT897272 |
| CY13 | Asia | China | Yuanjiang | CY | YJ13 | rice | NJAU | MT897273 |
| CY14 | Asia | China | Yuanjiang | CY | YJ14 | rice | NJAU | MT897274 |
| CX01 | Asia | China | Xinping | CX | XP1 | rice | NJAU | MT897275 |
| CX02 | Asia | China | Xinping | CX | XP2 | rice | NJAU | MT897276 |
| CX03 | Asia | China | Xinping | CX | XP3 | rice | NJAU | MT897277 |
| CX04 | Asia | China | Xinping | CX | XP4 | rice | NJAU | MT897278 |
| CX05 | Asia | China | Xinping | CX | XP5 | rice | NJAU | MT897279 |
| CX06 | Asia | China | Xinping | CX | XP6 | rice | NJAU | MT897280 |
| CX07 | Asia | China | Xinping | CX | XP7 | rice | NJAU | MT897281 |
| CX08 | Asia | China | Xinping | CX | XP8 | rice | NJAU | MT897282 |
| CX09 | Asia | China | Xinping | CX | XP9 | rice | NJAU | MT897283 |
| CX10 | Asia | China | Xinping | CX | XP10 | rice | NJAU | MT897284 |
| CX11 | Asia | China | Xinping | CX | XP11 | rice | NJAU | MT897285 |
| CX12 | Asia | China | Xinping | CX | XP12 | rice | NJAU | MT897286 |
| CX13 | Asia | China | Xinping | CX | XP13 | rice | NJAU | MT897287 |
| CX14 | Asia | China | Xinping | CX | XP14 | rice | NJAU | MT897288 |
| CX15 | Asia | China | Xinping | CX | XP15 | rice | NJAU | MT897289 |
| CC01 | Asia | China | Cangyuan | CC | CY1 | rice | NJAU | MT897290 |
| CC02 | Asia | China | Cangyuan | CC | CY2 | rice | NJAU | MT897291 |
| CC03 | Asia | China | Cangyuan | CC | CY3 | rice | NJAU | MT897292 |
| CC04 | Asia | China | Cangyuan | CC | CY4 | rice | NJAU | MT897293 |
| CC05 | Asia | China | Cangyuan | CC | CY5 | rice | NJAU | MT897294 |
| CC06 | Asia | China | Cangyuan | CC | CY6 | rice | NJAU | MT897295 |
| CC07 | Asia | China | Cangyuan | CC | CY7 | rice | NJAU | MT897296 |
| CC08 | Asia | China | Cangyuan | CC | CY8 | rice | NJAU | MT897297 |
| CC09 | Asia | China | Cangyuan | CC | CY9 | rice | NJAU | MT897298 |
| CC10 | Asia | China | Cangyuan | CC | CY10 | rice | NJAU | MT897299 |
| CC11 | Asia | China | Cangyuan | CC | CY11 | rice | NJAU | MT897300 |
| CC12 | Asia | China | Cangyuan | CC | CY12 | rice | NJAU | MT897301 |
| CC13 | Asia | China | Cangyuan | CC | CY13 | rice | NJAU | MT897302 |

| | | | | | | | | |
|---|---|---|---|---|---|---|---|---|
| CC15 | Asia | China | Cangyuan | CC | CY15 | rice | NJAU | MT897303 |
| CC16 | Asia | China | Cangyuan | CC | CY16 | rice | NJAU | MT897304 |
| CC17 | Asia | China | Cangyuan | CC | CY17 | rice | NJAU | MT897305 |
| CC18 | Asia | China | Cangyuan | CC | CY18 | rice | NJAU | MT897306 |
| CC19 | Asia | China | Cangyuan | CC | CY19 | rice | NJAU | MT897307 |
| CC20 | Asia | China | Cangyuan | CC | CY20 | rice | NJAU | MT897308 |
| IN01 | Asia | India | | IN | B1 | rice | INRAE DGIMI | MT897309 |
| IN02 | Asia | India | | IN | B2 | rice | INRAE DGIMI | MT897310 |
| IN03 | Asia | India | | IN | B3 | rice | INRAE DGIMI | MT897311 |
| IN05 | Asia | India | | IN | B5 | rice | INRAE DGIMI | MT897312 |
| IN06 | Asia | India | | IN | B6 | rice | INRAE DGIMI | MT897313 |
| IN07 | Asia | India | | IN | B7 | rice | INRAE DGIMI | MT897314 |
| IN08 | Asia | India | | IN | B8 | rice | INRAE DGIMI | MT897315 |
| IN09 | Asia | India | | IN | F1a | rice | INRAE DGIMI | MT897316 |
| IN10 | Asia | India | | IN | B10 | rice | INRAE DGIMI | MT897317 |
| IN11 | Asia | India | | IN | M1a | rice | INRAE DGIMI | MT897318 |
| IN12 | Asia | India | | IN | B12 | corn | INRAE DGIMI | MT897319 |
| IN13 | Asia | India | | IN | S3 | rice | INRAE DGIMI | MT897320 |
| UG01 | Africa | Uganda | | UG | 142m | rice | CSIRO/NaCRRI | MT897321 |
| UG02 | Africa | Uganda | | UG | 174F | rice | CSIRO/NaCRRI | MT897322 |
| UG03 | Africa | Uganda | | UG | 293m | corn | CSIRO/NaCRRI | MT897323 |
| UG04 | Africa | Uganda | | UG | C1 | rice | CSIRO/NaCRRI | MT897324 |
| UG05 | Africa | Uganda | | UG | KAB_48_2_S | rice | CSIRO/NaCRRI | MT897325 |
| UG06 | Africa | Uganda | | UG | KAT_40_19_S | corn | CSIRO/NaCRRI | MT897326 |
| UG07 | Africa | Uganda | | UG | KAT_37 | rice | CSIRO/NaCRRI | MT897327 |
| UG08 | Africa | Uganda | | UG | KAT_47 | rice | CSIRO/NaCRRI | MT897328 |
| UG09 | Africa | Uganda | | UG | KUM_7_27_S | rice | CSIRO/NaCRRI | MT897329 |
| UG10 | Africa | Uganda | | UG | NGO_16_8_S | rice | CSIRO/NaCRRI | MT897330 |
| UG11 | Africa | Uganda | | UG | RKT1 | rice | CSIRO/NaCRRI | MT897331 |
| UG12 | Africa | Uganda | | UG | SOR_29_48_S | rice | CSIRO/NaCRRI | MT897332 |
| UG13 | Africa | Uganda | | UG | STR1 | corn | CSIRO/NaCRRI | MT897333 |
| UG14 | Africa | Uganda | | UG | STR2 | rice | CSIRO/NaCRRI | MT897334 |
| UG15 | Africa | Uganda | | UG | STR3 | rice | CSIRO/NaCRRI | MT897335 |
| MW01 | Africa | Malawi | | MW | Mal1 | corn | CSIRO | MT897336 |
| MW02 | Africa | Malawi | | MW | Mal2 | rice | CSIRO | MT897337 |
| MW06 | Africa | Malawi | | MW | Mal6 | corn | CSIRO | MT897338 |
| MW07 | Africa | Malawi | | MW | Mal7 | corn | CSIRO | MT897339 |
| MW12 | Africa | Malawi | | MW | Mal12 | rice | CSIRO | MT897340 |
| MW16 | Africa | Malawi | | MW | Mal16 | corn | CSIRO | MT897341 |
| MW17 | Africa | Malawi | | MW | Mal17 | rice | CSIRO | MT897342 |
| MW21 | Africa | Malawi | | MW | Mal21 | rice | CSIRO | MT897343 |
| MW22 | Africa | Malawi | | MW | Mal22 | rice | CSIRO | MT897344 |
| MW26 | Africa | Malawi | | MW | Mal26 | corn | CSIRO | MT897345 |
| MW31 | Africa | Malawi | | MW | Mal31 | rice | CSIRO | MT897346 |
| MW32 | Africa | Malawi | | MW | Mal32 | corn | CSIRO | MT897347 |
| MW41 | Africa | Malawi | | MW | Mal41 | rice | CSIRO | MT897348 |
| MW42 | Africa | Malawi | | MW | Mal42 | rice | CSIRO | MT897349 |
| MW46 | Africa | Malawi | | MW | Mal46 | rice | CSIRO | MT897350 |
| MW51 | Africa | Malawi | | MW | Mal51 | rice | CSIRO | MT897351 |
| TZ01 | Africa | Tanzania | | TZ | 345K | rice | CSIRO | MT897352 |
| BE01 | Africa | Benin | | BE | BCW1 | rice | INRAE DGIMI | MT897353 |
| BE02 | Africa | Benin | | BE | BCW2 | rice | INRAE DGIMI | MT897354 |
| BE29 | Africa | Benin | | BE | KNAM_0029 | rice | INRAE DGIMI | MT897355 |
| BE30 | Africa | Benin | | BE | KNAM_0030 | corn | INRAE DGIMI | MT897356 |
| UF01 | NthAmerica | USA | Florida | UF | FCC1 | corn | INRAE DGIMI | MT897357 |
| UF02 | NthAmerica | USA | Florida | UF | FCC2 | corn | INRAE DGIMI | MT897358 |
| UF03 | NthAmerica | USA | Florida | UF | FCC3 | corn | INRAE DGIMI | MT897359 |
| UF04 | NthAmerica | USA | Florida | UF | FCC4 | corn | INRAE DGIMI | MT897360 |

| | | | | | | | | |
|---|---|---|---|---|---|---|---|---|
| UF05 | NthAmerica | USA | Florida | UF | FCC5 | corn | INRAE DGIMI | MT897361 |
| UF06 | NthAmerica | USA | Florida | UF | FCC6 | Corn | INRAE DGIMI | MT897362 |
| UF07 | NthAmerica | USA | Florida | UF | FCC7 | corn | INRAE DGIMI | MT897363 |
| UF08 | NthAmerica | USA | Florida | UF | FCC8 | corn | INRAE DGIMI | MT897364 |
| UF09 | NthAmerica | USA | Florida | UF | FGJ2 | rice | INRAE DGIMI | MT897365 |
| UF10 | NthAmerica | USA | Florida | UF | FGJ3 | rice | INRAE DGIMI | MT897366 |
| UF11 | NthAmerica | USA | Florida | UF | FGJ4 | corn | INRAE DGIMI | MT897367 |
| UF12 | NthAmerica | USA | Florida | UF | FGJ5 | rice | INRAE DGIMI | MT897368 |
| UF13 | NthAmerica | USA | Florida | UF | FGJ6 | rice | INRAE DGIMI | MT897369 |
| UF14 | NthAmerica | USA | Florida | UF | FGJ7 | rice | INRAE DGIMI | MT897370 |
| UF15 | NthAmerica | USA | Florida | UF | FGJ8 | rice | INRAE DGIMI | MT897371 |
| UF16 | NthAmerica | USA | Florida | UF | FGJ9 | rice | INRAE DGIMI | MT897372 |
| UF17 | NthAmerica | USA | Florida | UF | FGJ10 | rice | INRAE DGIMI | MT897373 |
| UF18 | NthAmerica | USA | Florida | UF | FGJ11 | rice | INRAE DGIMI | MT897374 |
| UF19 | NthAmerica | USA | Florida | UF | FGJ12 | rice | INRAE DGIMI | MT897375 |
| UF20 | NthAmerica | USA | Florida | UF | FL16 | corn | INRAE DGIMI | MT897376 |
| UF21 | NthAmerica | USA | Florida | UF | FL17 | corn | INRAE DGIMI | MT897377 |
| UF22 | NthAmerica | USA | Florida | UF | FL18 | corn | INRAE DGIMI | MT897378 |
| UF23 | NthAmerica | USA | Florida | UF | FL19 | corn | INRAE DGIMI | MT897379 |
| UF24 | NthAmerica | USA | Florida | UF | FL20 | corn | INRAE DGIMI | MT897380 |
| UM01 | NthAmerica | USA | Mississippi | UM | MS_C_C1 | corn | INRAE DGIMI | MT897381 |
| UM02 | NthAmerica | USA | Mississippi | UM | MS_C_C2 | corn | INRAE DGIMI | MT897382 |
| UM03 | NthAmerica | USA | Mississippi | UM | MS_C_C3 | corn | INRAE DGIMI | MT897383 |
| UM04 | NthAmerica | USA | Mississippi | UM | MS_C_C4 | corn | INRAE DGIMI | MT897384 |
| UM05 | NthAmerica | USA | Mississippi | UM | MS_C_C5 | corn | INRAE DGIMI | MT897385 |
| UM06 | NthAmerica | USA | Mississippi | UM | MS_C_C6 | corn | INRAE DGIMI | MT897386 |
| UM07 | NthAmerica | USA | Mississippi | UM | MS_C_C7 | corn | INRAE DGIMI | MT897387 |
| UM08 | NthAmerica | USA | Mississippi | UM | MS_C_C8 | corn | INRAE DGIMI | MT897388 |
| UM09 | NthAmerica | USA | Mississippi | UM | MS_C_C9 | corn | INRAE DGIMI | MT897389 |
| UM10 | NthAmerica | USA | Mississippi | UM | MS_R_R1 | rice | INRAE DGIMI | MT897390 |
| UM11 | NthAmerica | USA | Mississippi | UM | MS_R_R2 | rice | INRAE DGIMI | MT897391 |
| UM12 | NthAmerica | USA | Mississippi | UM | MS_R_R3 | rice | INRAE DGIMI | MT897392 |
| UM13 | NthAmerica | USA | Mississippi | UM | MS_R_R4 | rice | INRAE DGIMI | MT897393 |
| UM14 | NthAmerica | USA | Mississippi | UM | MS_R_R5 | rice | INRAE DGIMI | MT897394 |
| UM15 | NthAmerica | USA | Mississippi | UM | MS_R_R6 | rice | INRAE DGIMI | MT897395 |
| UM16 | NthAmerica | USA | Mississippi | UM | MS_R_R7 | rice | INRAE DGIMI | MT897396 |
| UM17 | NthAmerica | USA | Mississippi | UM | MS_R_R8 | rice | INRAE DGIMI | MT897397 |
| UM18 | NthAmerica | USA | Mississippi | UM | MS_R_R9 | rice | INRAE DGIMI | MT897398 |
| PR01 | NthAmerica | Puerto-Rico | Puerto-Rico | PR | PR1 | corn | INRAE DGIMI | MT897399 |
| PR05 | NthAmerica | Puerto-Rico | Puerto-Rico | PR | PR5 | corn | INRAE DGIMI | MT897400 |
| PR12 | NthAmerica | Puerto-Rico | Puerto-Rico | PR | PR12 | corn | INRAE DGIMI | MT897401 |
| PR14 | NthAmerica | Puerto-Rico | Puerto-Rico | PR | PR14 | rice | INRAE DGIMI | MT897402 |
| PR15 | NthAmerica | Puerto-Rico | Puerto-Rico | PR | PR15 | corn | INRAE DGIMI | MT897403 |
| PR16 | NthAmerica | Puerto-Rico | Puerto-Rico | PR | PR16 | corn | INRAE DGIMI | MT897404 |
| PR18 | NthAmerica | Puerto-Rico | Puerto-Rico | PR | PR18 | corn | INRAE DGIMI | MT897405 |
| PR19 | NthAmerica | Puerto-Rico | Puerto-Rico | PR | PR19 | corn | INRAE DGIMI | MT897406 |
| PR27 | NthAmerica | Puerto-Rico | Puerto-Rico | PR | PR27 | rice | INRAE DGIMI | MT897407 |
| PR29 | NthAmerica | Puerto-Rico | Puerto-Rico | PR | PR29 | corn | INRAE DGIMI | MT897408 |
| PR30 | NthAmerica | Puerto-Rico | Puerto-Rico | PR | PR30 | rice | INRAE DGIMI | MT897409 |
| PR31 | NthAmerica | Puerto-Rico | Puerto-Rico | PR | PR31 | corn | INRAE DGIMI | MT897410 |
| PR32 | NthAmerica | Puerto-Rico | Puerto-Rico | PR | PR32 | corn | INRAE DGIMI | MT897411 |
| PR33 | NthAmerica | Puerto-Rico | Puerto-Rico | PR | PR33 | corn | INRAE DGIMI | MT897412 |
| PR35 | NthAmerica | Puerto-Rico | Puerto-Rico | PR | PR35 | rice | INRAE DGIMI | MT897413 |
| MX39 | NthAmerica | Mexico | | MX | KNAM_0039 | corn | INRAE DGIMI | MT897414 |
| MX40 | NthAmerica | Mexico | | MX | KNAM_0040 | corn | INRAE DGIMI | MT897415 |
| MX41 | NthAmerica | Mexico | | MX | KNAM_0041 | corn | INRAE DGIMI | MT897416 |
| MX42 | NthAmerica | Mexico | | MX | KNAM_0042 | corn | INRAE DGIMI | MT897417 |
| MX43 | NthAmerica | Mexico | | MX | KNAM_0043 | corn | INRAE DGIMI | MT897418 |

| | | | | | | | | |
|---|---|---|---|---|---|---|---|---|
| MX44 | NthAmerica | Mexico | | MX | KNAM_0044 | corn | INRAE DGIMI | MT897419 |
| MX45 | NthAmerica | Mexico | | MX | KNAM_0045 | corn | INRAE DGIMI | MT897420 |
| MX46 | NthAmerica | Mexico | | MX | KNAM_0046 | corn | INRAE DGIMI | MT897421 |
| MX47 | NthAmerica | Mexico | | MX | KNAM_0047 | corn | INRAE DGIMI | MT897422 |
| MX48 | NthAmerica | Mexico | | MX | KNAM_0048 | corn | INRAE DGIMI | MT897423 |
| GP01 | SthAmerica | Guadaloup | | GP | G_1.1 | rice | INRAE DGIMI | MT897424 |
| GP02 | SthAmerica | Guadaloup | | GP | G_1.2 | rice | INRAE DGIMI | MT897425 |
| GP03 | SthAmerica | Guadaloup | | GP | G_2.2 | rice | INRAE DGIMI | MT897426 |
| GP04 | SthAmerica | Guadaloup | | GP | G_2.3 | rice | INRAE DGIMI | MT897427 |
| GF02 | SthAmerica | French-Guiana | | GF | FG2 | rice | INRAE DGIMI | MT897428 |
| GF03 | SthAmerica | French-Guiana | | GF | FG3 | rice | INRAE DGIMI | MT897429 |
| GF04 | SthAmerica | French-Guiana | | GF | FG4 | rice | INRAE DGIMI | MT897430 |
| PE01 | SthAmerica | Peru | | PE | HIA242 | corn | CSIRO/DAWE | MT897431 |
| PE02 | SthAmerica | Peru | | PE | HIA263 | corn | CSIRO/DAWE | MT897432 |
| PE03 | SthAmerica | Peru | | PE | HIA273 | corn | CSIRO/DAWE | MT897433 |
| PE04 | SthAmerica | Peru | | PE | HIA309 | corn | CSIRO/DAWE | MT897434 |
| PE05 | SthAmerica | Peru | | PE | HIA312 | corn | CSIRO/DAWE | MT897435 |
| PE06 | SthAmerica | Peru | | PE | HIA32 | corn | CSIRO/DAWE | MT897436 |
| PE07 | SthAmerica | Peru | | PE | HIA33 | corn | CSIRO/DAWE | MT897437 |
| PE08 | SthAmerica | Peru | | PE | HIA44 | corn | CSIRO/DAWE | MT897438 |
| PE09 | SthAmerica | Peru | | PE | HIA52 | corn | CSIRO/DAWE | MT897439 |
| PE10 | SthAmerica | Peru | | PE | HIA53 | corn | CSIRO/DAWE | MT897440 |
| PE11 | SthAmerica | Peru | | PE | HIA7 | corn | CSIRO/DAWE | MT897441 |
| PE12 | SthAmerica | Peru | | PE | HIA74 | corn | CSIRO/DAWE | MT897442 |
| PE13 | SthAmerica | Peru | | PE | HIA75 | corn | CSIRO/DAWE | MT897443 |
| PE14 | SthAmerica | Peru | | PE | HIA8 | corn | CSIRO/DAWE | MT897444 |
| PE15 | SthAmerica | Peru | | PE | HIA90 | corn | CSIRO/DAWE | MT897445 |
| PE16 | SthAmerica | Peru | | PE | HIA98 | corn | CSIRO/DAWE | MT897446 |
| BC38 | SthAmerica | Brazil | | BC | CC38 | corn | CSIRO/UFG | MT897447 |
| BC40 | SthAmerica | Brazil | | BC | CC40 | corn | CSIRO/UFG | MT897448 |
| BC41 | SthAmerica | Brazil | | BC | CC41 | corn | CSIRO/UFG | MT897449 |
| BC46 | SthAmerica | Brazil | | BC | CC46 | rice | CSIRO/UFG | MT897450 |
| BC49 | SthAmerica | Brazil | | BC | CC49 | corn | CSIRO/UFG | MT897451 |
| BC55 | SthAmerica | Brazil | | BC | CC55 | corn | CSIRO/UFG | MT897452 |
| BC61 | SthAmerica | Brazil | | BC | CC61 | corn | CSIRO/UFG | MT897453 |
| BC66 | SthAmerica | Brazil | | BC | CC66 | corn | CSIRO/UFG | MT897454 |
| BR01 | SthAmerica | Brazil | | BR | rCC1 | corn | CSIRO/UFG | MT897455 |
| BR05 | SthAmerica | Brazil | | BR | rCC5 | corn | CSIRO/UFG | MT897456 |
| BR20 | SthAmerica | Brazil | | BR | rCC20 | corn | CSIRO/UFG | MT897457 |
| BR25 | SthAmerica | Brazil | | BR | rCC25 | corn | CSIRO/UFG | MT897458 |

**Supplemental Table 2:** Neutrality test statistics for *Spodoptera frugiperda* (FAW) from native (Brazil, French Guina, Guadeloupe, Mexico, Peru, Puerto Rico, USA) and invasive (Benin, Cina, India, Malawi, Tanzania, Uganda) ranges. Estimates are based on 870 genome-wide single nucleotide polymorphic markers (see Material & Methods) and are not made for populations with ≤ 3 individuals (CH, GF, TZ).

| Pop. Code | Pop. | No. Samples | Tajima's D | Fu & Li's D* |
|---|---|---|---|---|
| BC | Brazil-CC | 8 | 0.693 | 0.671 |
| BE | Benin | 4 | 0.331 | 0.328 |
| BR | Brazil-rCC | 4 | 0.13 | 0.13 |
| CC | China-CY | 19 | 1.418 | 1.286 |
| CH | China-H06 | 1 | | |
| CX | China-XP | 15 | 1.249 | 1 |
| CY | China-YJ | 12 | 1.013 | 1 |
| GF | French Guiana | 3 | | |
| GP | Guadeloupe | 4 | 0.293 | 0 |
| IN | India | 12 | 1.137 | 1 |
| MW | Malawi | 16 | 1.411 | 1.3 |
| MX | Mexico | 10 | 0.717 | 0.61 |
| PE | Peru | 16 | 1.554 | 1.293 |
| PR | Puerto Rico | 15 | 1.241 | 1.188 |
| TZ | Tanzania | 1 | | |
| UF | USA-FL | 24 | 1.47 | 1.699 |
| UG | Uganda | 15 | 1.795 | 1.298 |
| UM | USA-MS | 18 | 1.717 | 1.452 |

**Note:** The statistical tests for neutrality were highest for populations with highest $H_{obs}$ and lowest numbers of loci in HWE. For the Tajima's D estimates, all populations showed positive D values, particularly for most of the invasive populations (especially values > 1.40) but also native populations from Florida, Mississippi, and Peru, suggesting either there was a lack of rare alleles in these populations; that these populations experienced sudden contraction; or that balancing selection was acting to maintain heterozygotes (e.g., due to heterozygous advantage). Lower Tajima's D values were detected in native (e.g., Brazil-rCC, Guadeloupe) and invasive (e.g., Benin) populations, suggesting that these were evolving at close to mutation-drift equilibrium. The other test for neutrality, Fu & Li's D*, gave consistent results with Tajima's D to support that particularly in invasive populations, there was a lack of singleton alleles that represented either a recent bottleneck, population reduction, population subdivision, or migration, rather than that these invasive populations have undergone recent expansion since the western African bottleneck/founder event. Note that the high Tajima D and Fu & Li's D* estimates from this study should be considered with caution due to the uneven and at times very low populatin sample sizes as well as high number of makers used, with nuetrality tests such as Tajima's D and Fu & Li's D* estimates being known to be impacted by sample sizes[1]
